# Tribbles1 and Cop1 cooperate to protect the host during *in vivo* mycobacterial infection

**DOI:** 10.1101/2023.08.25.553505

**Authors:** Ffion R Hammond, Amy Lewis, Gabriele Pollara, Gillian S Tomlinson, Mahdad Noursadeghi, Endre Kiss-Toth, Philip M Elks

## Abstract

Tuberculosis is a major global health problem and is one of the top 10 causes of death worldwide. There is a pressing need for new treatments that circumvent emerging antibiotic resistance. *Mycobacterium tuberculosis* parasitises macrophages, reprogramming them to establish a niche in which to proliferate, therefore macrophage manipulation is a potential host-directed therapy if druggable molecular targets could be identified. The pseudokinase Tribbles1 (Trib1) regulates multiple innate immune processes and inflammatory profiles making it a potential drug target in infections. Trib1 controls macrophage function, cytokine production and macrophage polarisation. Despite wide-ranging effects on leukocyte biology, data exploring the roles of Tribbles in infection *in vivo* are limited. Here, we identify that human Tribbles 1 is expressed in monocytes and is upregulated at the transcript level after stimulation with mycobacterial antigen. To investigate the mechanistic roles of Tribbles in the host response to mycobacteria *in vivo*, we used a zebrafish *Mycobacterium marinum* (Mm) infection tuberculosis model. Zebrafish Tribbles family members were characterised and shown to have substantial mRNA and protein sequence homology to their human orthologues. *trib1* overexpression was host-protective against Mm infection, reducing burden by approximately 50%. Conversely, *trib1* knockdown exhibited increased infection. Mechanistically, *trib1* overexpression significantly increased the levels of pro-inflammatory factors *il-1*β and nitric oxide. The host-protective effect of *trib1* was found to be dependent on the E3 ubiquitin kinase Cop1. These findings highlight the importance of Trib1 and Cop1 as immune regulators during infection *in vivo* and suggest that enhancing macrophage TRIB1 levels may provide a tractable therapeutic intervention to improve bacterial infection outcomes in tuberculosis.

## Introduction

With the rise of anti-microbial resistance (AMR), bacterial infections are a major threat to global public health. Tuberculosis, caused by the human pathogen *Mycobacterium tuberculosis*, is a case in point, with 1.6 million deaths worldwide (WHO 2022), many of which are resistant to first- and second-line antibiotic treatments (Allué-Guardia et al. 2021; Hameed et al. 2018; Migliori et al. 2013). To successfully combat AMR there is a pressing and urgent need for alternative treatment strategies to failing antimicrobials. One such approach is offered by the development of host derived therapies (HDT), which target systems in the host rather than the pathogen, circumventing AMR (Kaufmann et al. 2018; Kilinç et al. 2021).

One primary immune defence against *Mycobacterium tuberculosis* (*Mtb*) is macrophages. Macrophages have a spectrum of phenotypes ranging from proinflammatory to anti-inflammatory, determined in a process known as macrophage polarisation. *Mtb* is expert at manipulation of macrophage polarisation to its advantage (Ahmad et al. 2022) and can inhibit the polarisation of proinflammatory macrophages, subverting killing mechanisms to promote intracellular survival of the bacteria and subsequent granuloma formation (Hackett et al. 2020). Reprogramming macrophages to better kill *Mtb* is a potential HDT strategy that may be particularly effective against intracellular pathogens (Sheedy and Divangahi 2021).

Tribbles genes encode for a family of pseudokinases (TRIB1, TRIB2 and TRIB3 (Kiss-Toth et al. 2004)), involved in the regulation of core cellular processes, ranging from cell cycle to glucose metabolism (Grosshans and Wieschaus 2000; Mata et al. 2000; Seher and Leptin 2000). The TRIB1 isoform has been strongly associated with macrophage roles in inflammation and innate immunity (Johnston et al. 2015; Niespolo et al. 2020)). TRIB1 regulates multiple important macrophage regulatory factors, especially controlling the proinflammatory response, such as tumour necrosis factor alpha (TNF-α), interleukin-1beta (IL-1β) and nitric oxide (NO) (Arndt et al. 2018; Liu et al. 2013). *Trib1^-/-^* mice have decreased expression levels of inflammation related genes such as *IL-6*, *IL-1b* and *Nos2* (encodes for inducible nitric oxide synthase, iNOS), and murine *Trib1-/-* bone marrow derived macrophages have defective inflammatory, phagocytic, migratory and NO responses *in vitro* (Arndt et al. 2018; Liu et al. 2013).

TRIB1 influences inflammatory and immune processes via multiple mechanisms. The best described is via recruitment and binding of the E3 ubiquitin ligase constitutive photomorphogenic 1 (COP1). The TRIB1 protein possesses two functional binding sites in its C-terminal, one for constitutive photomorphogenic 1 (COP1) and the second for Mitogen-activated protein kinase kinase (MEK) binding. TRIB1 can act as a protein scaffold, binding a substrate to its pseudokinase domain, as well as binding in the functional C terminus to create a regulatory complex. Binding of TRIB1 to the E3 ubiquitin ligase COP1 causes a conformational change, enhancing COP1 binding and bringing COP1 into proximity with the substrate allowing ubiquitination and subsequent degradation (Jamieson et al. 2018; Kung and Jura 2019; Murphy et al. 2015; Zahid et al. 2022). The TRIB1/COP1 complex is responsible for the regulation of multiple targets such as transcription factors, including the tumour suppressor CCAAT/enhancer-binding protein (C/EBPα), which regulates macrophage migration and TNF-α production (Liu et al. 2013; Yoshida et al. 2013).

While TRIB1 has been shown to regulate several inflammatory and innate immune functions *in vitro*, its role in infection is much less characterised, especially in an *in vivo* setting. *TRIB1* is a predicted target of microRNA-gene interactions that differentiate active and latent TB patients (Wu et al. 2014) and is an overabundant transcript in highly pro-inflammatory tuberculosis-immune reconstitution inflammatory syndrome (TB-IRIS) patients (Lai et al. 2015). However, despite these reported potential links between Tribbles and TB, interrogation of *TRIB* isoform transcripts in human mycobacterial datasets had not been performed.

Over the last two decades, the zebrafish has proved a powerful model for understanding host-pathogen interactions, due to its high-fecundity, transparency of larvae and availability of transgenic reporter lines. A human disease-relevant and tractable infection model is the zebrafish model of tuberculosis, utilising the injection of the natural fish pathogen *Mycobacterium marinum* (*Mm*), a close genetic relative of *<u>Mtb</u>*,(Davis et al. 2002; van der Sar et al. 2009). This model has shed light on numerous immune pathways involved in host defence, for example Hypoxia Inducible Factor (HIF) signalling (Elks et al. 2013; Ogryzko et al. 2019; Schild et al. 2020).

Here, we show that Tribbles 1 is expressed in human primary monocytes and its expression is increased at the site of a human *in vivo* mycobacterial antigen challenge, indicative of a role in TB responses. To substantiate the importance of *TRIB1* in TB pathogenesis, we report a new, protective role for *trib1* in infection defence using an *in vivo* zebrafish *Mycobacterium marinum* (Mm) infection model. Overexpression of *trib1* significantly reduced Mm burden and increased production of the pro-inflammatory cytokine *il1b* and NO. The antimicrobial effect of *trib1* overexpression was found to be dependent on *cop1*. Our findings uncover a role for *trib1* in mycobacterial infection defence *in vivo*, highlighting Trib1 as a potential therapeutic target for manipulation to improve bacterial infection outcomes.

## Materials and Methods

### Human transcriptomic dataset analysis

Expression of TRIB1 in human CD14+ monocytes and the site of a tuberculin skin test (TST) was derived from publicly available transcriptomic data deposited in EBI ArrayExpress repository (datasets E-MTAB-8162 & E-MTAB-6816 respectively - https://www.ebi.ac.uk/biostudies/arrayexpress) (Pollara et al. 2021).

### Zebrafish

Zebrafish were raised in The Biological Services Aquarium (University of Sheffield, UK) and maintained according to standard protocols (zfin.org) in Home Office approved facilities. All procedures were performed on embryos pre 5.2 days post fertilisation (dpf) which were therefore outside of the Animals (Scientific Procedures) Act, to standards set by the UK Home Office. Adult fish were maintained at 28°C with a 14/10-hour light/dark cycle. Nacre zebrafish were used as a wildtype. Transgenic zebrafish lines used are detailed below in Table 1.

**Table 1:**
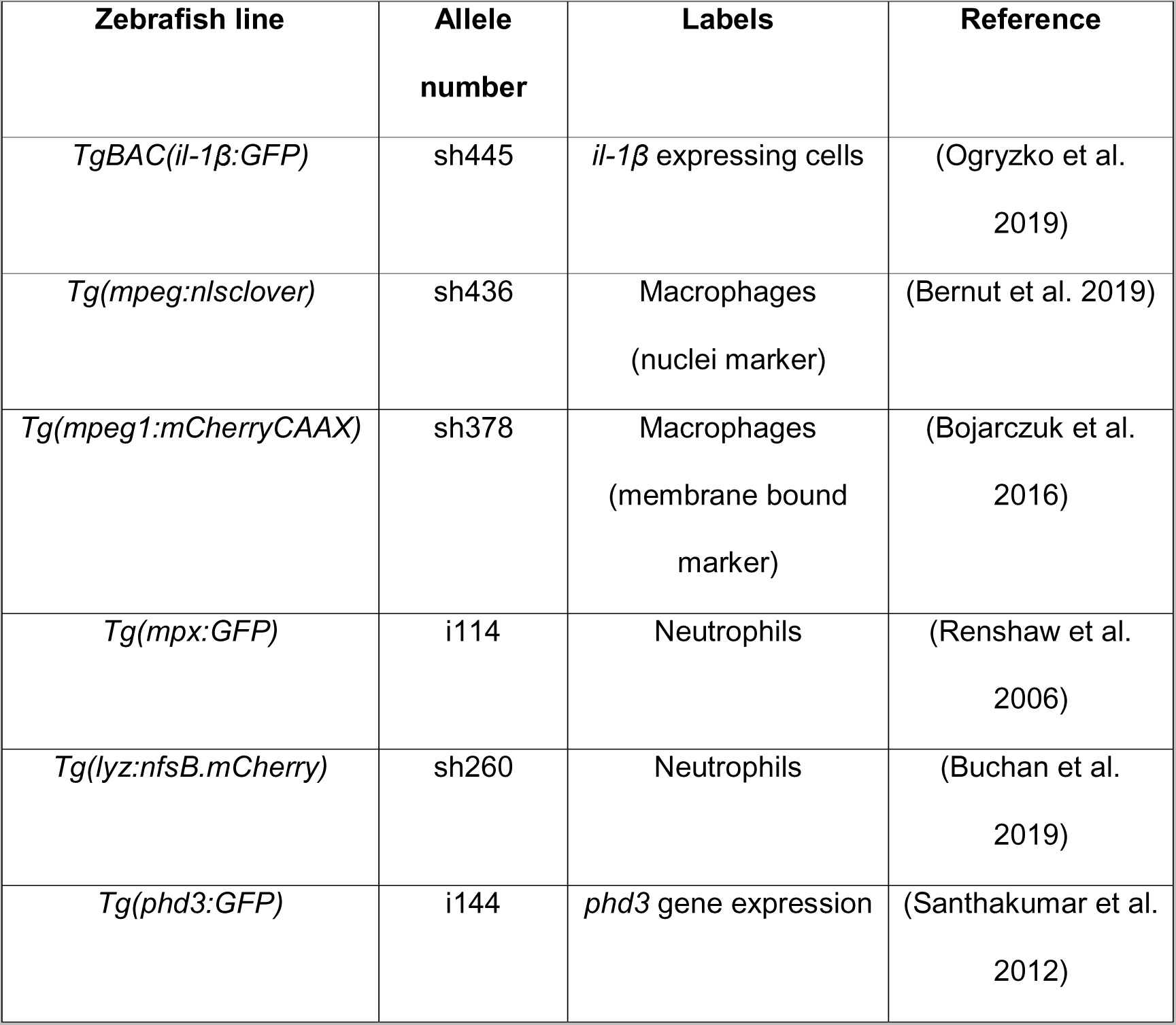
Transgenic zebrafish lines.

### CRISPR-Cas9 guide design and CRISPant generation

Transcript details for *trib1* (current Ensembl entry code is ENSDARG00000110963, but previously coded as ENSDARG00000076142 which is the identifier code used in RNAseq datasets), *trib2* (ENSDARG00000068179) and *trib3* (ENSDARG00000016200) were obtained from Ensembl genome browser (www.ensembl.org). Only one transcript was identified per gene which was used for CRISPR-Cas9 guide design. The web tool ChopChop (https://chopchop.cbu.uib.no) was used to design guideRNAs and primers. A summary of all guideRNAs (Sigma-Aldrich) and primer oligos (IDT) designed is described in Table 2 below.

**Table 2:**
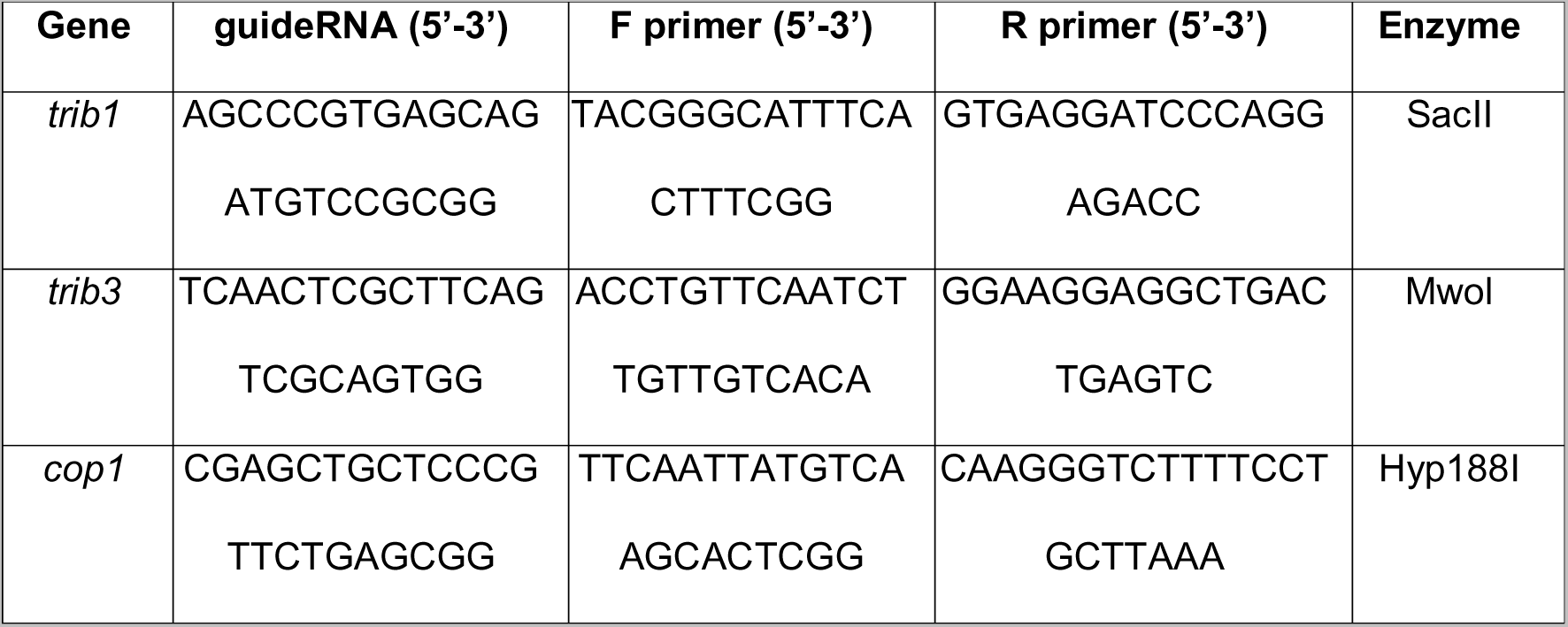
Summary of CRISPR-Cas9 guideRNAs, relevant primers and restriction enzymes used for genotyping.

To genotype first genomic DNA was extracted from 2-4dpf larvae via incubation at 95°C in 100µl of 50mM NaOH for 20 minutes followed by the addition of 10µl 1M Tris-HCl (pH8). PCR was then performed on genomic DNA with relevant primer pair and enzyme (NEB) (see materials and methods chapter for PCR programme). Digests were run on a 2% (w/v) agarose gel (Appleton Woods) at 100v. Samples that were positive for CRISPR mutation were not digested by the restriction enzyme due to destruction of the restriction enzyme recognition site.

All guideRNAs (Sigma/Merck) were microinjected in the following injection mix: 1μl 20mM guideRNA, 1μl 20mM Tracr RNA (Sigma/Merck), 1μl Cas9 (diluted 1:3 in diluent B, NEB), 1μl water (water was replaced with 100ng/μl *trib1* RNA for *cop1* experiments). A *tyrosinase* guideRNA (Sigma/Merck) control that has negligible effects on innate immunity was used as a negative CRISPR (Isles et al. 2019). Embryos were microinjected with 1nl guideRNA mix at the single cell stage to generate F0 CRISPants.

### Cloning and whole mount in situ hybridisation of trib 1, 2 and 3

RNA probes for zebrafish *trib1* (ENSDARG00000110963), *trib2* (ENSDARG00000068179) and *trib3* (ENSDARG00000016200) were designed and synthesised after cloning the full-length genes into the pCR™Blunt II-TOPO® vector according to manufacturer’s instructions (ThermoFisher Scientific). Plasmid was linearised with the relevant restriction enzyme (Table 3), NEB Biolabs) and probes were synthesised according to DIG RNA Labelling Kit (SP6/T7, Roche). Zebrafish larvae were anaesthetised in 0.168mg/ml Tricaine (MS-222, Sigma-Aldrich) in E3 media, which was removed and replaced with 4% (v/v in PBS) paraformaldehyde solution (PFA, ThermoFisher Scientific) overnight at 4°C to fix. Whole mount *in situ* hybridisation was performed as previously described (Thisse and Thisse 2008).

**Table 3:**
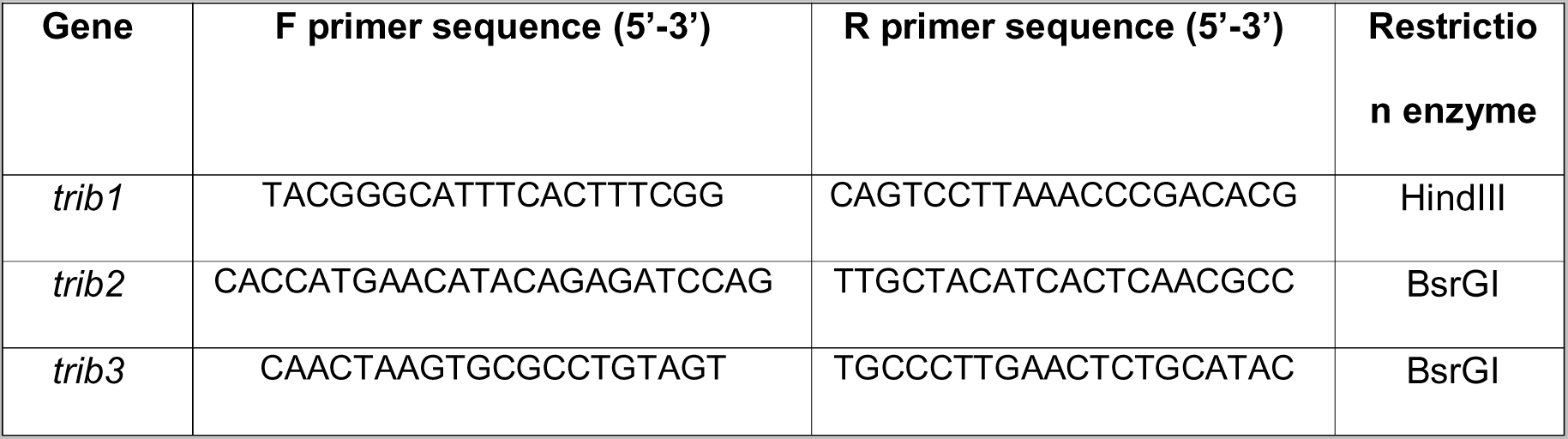
Primers used for tribbles PCR for TOPO transformation and relevant restriction enzymes used for linearisation.

### RNA injections for trib overexpression experiments

Forward inserts of *trib1*, *trib2* and *trib3* were cut from the pCR™Blunt II-TOPO® constructs using a double restriction digest with BamHI and XbaI at 37°C for 1.5 hours. The expression vector pCS2+ (Addgene) was digested using the same restriction enzyme pair and all digests were gel extracted using QIAquick Gel Extraction Kit (Qiagen). Gel extracts of vector and *trib* digests were ligated into pCS2+ via overnight incubation at room temperature with T4 DNA ligase according to manufacturer’s instructions (NEB). Constructs were confirmed using sequencing performed by the University of Sheffield’s Genomics core facility. RNA of each *trib* isoform was transcribed using mMessageMachine kit (Ambion, Invitrogen) and diluted to 100ng/μl in phenol red (PR, diluted 1:10 in RNAse free water) for microinjection. Embryos were microinjected with 1nl of 100ng/μl RNA (measured using a 10mm graticule) at the single cell stage as previously described (Elks et al. 2011). RNA of dominant active (DA) and negative (DN) *hif-1ab* variants (ZFIN: hif1ab) were used for controls (Elks et al. 2013; Elks et al. 2011).

### Mycobacterium marinum culture and injection

Bacterial infection experiments were performed using *Mycobacterium marinum* strain M (ATCC #BAA-535), containing the pSMT3-mCherry vector. Liquid cultures were prepared from bacterial plates before washing in PBS and diluting in 2% (w/v) polyvinylpyrrolidone40 (PVP40, Sigma-Aldrich) for injection as described previously (Benard et al. 2012). Injection inoculum was prepared to 100 colony forming units (cfu)/nl for all burden experiments, loaded into borosilicate glass microcapillary injection needles (WPI, pulled using a micropipette puller device, WPI) before microinjection into the circulation of 30hpf zebrafish larvae via the caudal vein.

Prior to injection, zebrafish were anaesthetised in 0.168 mg/ml Tricaine (MS-222, Sigma-Aldrich) in E3 media and were transferred onto 1% agarose in E3+methylene blue plates, removing excess media. All pathogens were injected using a microinjection rig (WPI) attached to a dissecting microscope. A 10mm graticule was used to measure 1nl droplets of injection volume, and for consistency, droplets were tested every 5-10 fish and the needle recalibrated if necessary. After injection, zebrafish were transferred to fresh E3 media for recovery and maintained at 28°C.

### Anti-nitrotyrosine immunostaining

Larvae were fixed in 4% (v/v) paraformaldehyde in PBS overnight at 4°C, and nitrotyrosine levels were immune labelled using immunostaining with a rabbit polyclonal anti-nitrotyrosine antibody (06-284; Merck Millipore) and detected using an Alexa Fluor–conjugated secondary antibody (Invitrogen Life Technologies) as previously described (Elks et al. 2014; Elks et al. 2013).

### Confocal microscopy

*TgBAC(il-1*β*:GFP)sh445* larvae and larvae immune-stained for nitrotyrosine were imaged using a Leica DMi8 SPE-TCS microscope using a HCX PL APO 40x/1,10 water immersion lens. Larvae were anaesthetised in 0.168 mg/ml Tricaine and mounted in 1% (w/v) low melting agarose (Sigma) containing 0.168 mg/ml tricaine (Sigma) in 15μ-Slide 4 well glass bottom slides (Ibidi).

### Stereo microscopy

Zebrafish larvae were anaesthetised in 0.168 mg/ml Tricaine and transferred to a 50mm glass bottomed FluoroDish^TM^ (Ibidi). Zebrafish were imaged using a Leica DMi8 SPE-TCS microscope fitted with a Hamamatsu ORCA Flash 4.0 camera attachment using a HC FL PLAN 2.5x/0.07 dry lens. Whole mount *in situ* staining was imaged using a Leica MZ10F stereo 14 microscope fitted with a GXCAM-U3 series 5MP camera (GT Vision).

### Image analysis

To calculate bacterial burden, fluorescent pixel count was measured using dedicated pixel count software (Stoop et al. 2011). For confocal imaging of anti-nitrotyrosine staining or transgenic lines, ImageJ (Schindelin et al. 2012) was used to quantify corrected total cell fluorescence (CTCF) (Elks et al. 2014; Elks et al. 2013).

### Statistical analysis

Statistical significance was calculated and determined using Graphpad Prism 9.0. Quantified data figures display all datapoints, with error bars depicting standard error of the mean (SEM) unless stated otherwise in the figure legend. Statistical significance was determined using one-way ANOVA with Bonferroni’s multiple comparisons post hoc test/Kruskal Wallis for experiments with three or more experimental groups, or paired/unpaired T test/Wilcoxon matched pairs signed rank test for experiments with two experimental groups, unless stated otherwise in figure legend. P values shown are: \**P* < .05, \*\**P* < .01, and \*\*\**P* < .001.

## Results

### *TRIB1* is expressed in human monocytes and is upregulated after *in vivo* mycobacterial antigen stimulation

To explore whether Tribble pseudokinase expression is modulated by mycobacterial antigen exposure in humans, we initially focused on CD14+ monocytes stimulated *in vitro* with *Mtb* protein derivative (PPD). This revealed that mycobacterial antigen exposure induced the expression of TRIB1 isoform transcripts but not TRIB2, which had the lowest baseline expression, nor TRIB3, observations consistent across monocytes from either active or latent TB individuals (Figure 1A-C). To determine whether Tribbles play a role in human responses *in vivo*, we turned to the transcriptomic profiles of biopsies from the site of a tuberculin skin test (TST), a routine clinical investigation repurposed into a mycobacterial antigen challenge model (Bell et al. 2016). This revealed baseline expression of TRIB 1 in control saline injected tissue samples (Figure 1E). Exposure to tuberculin induced robust induction of TRIB1 expression in TST reactions for both active and latent TB individuals (Figure 1E). A more modest increase was seen for TRIB2 (Figure 1F) but not for TRIB3 (Figure 1G) (Pollara et al. 2021).

**Figure 1:**
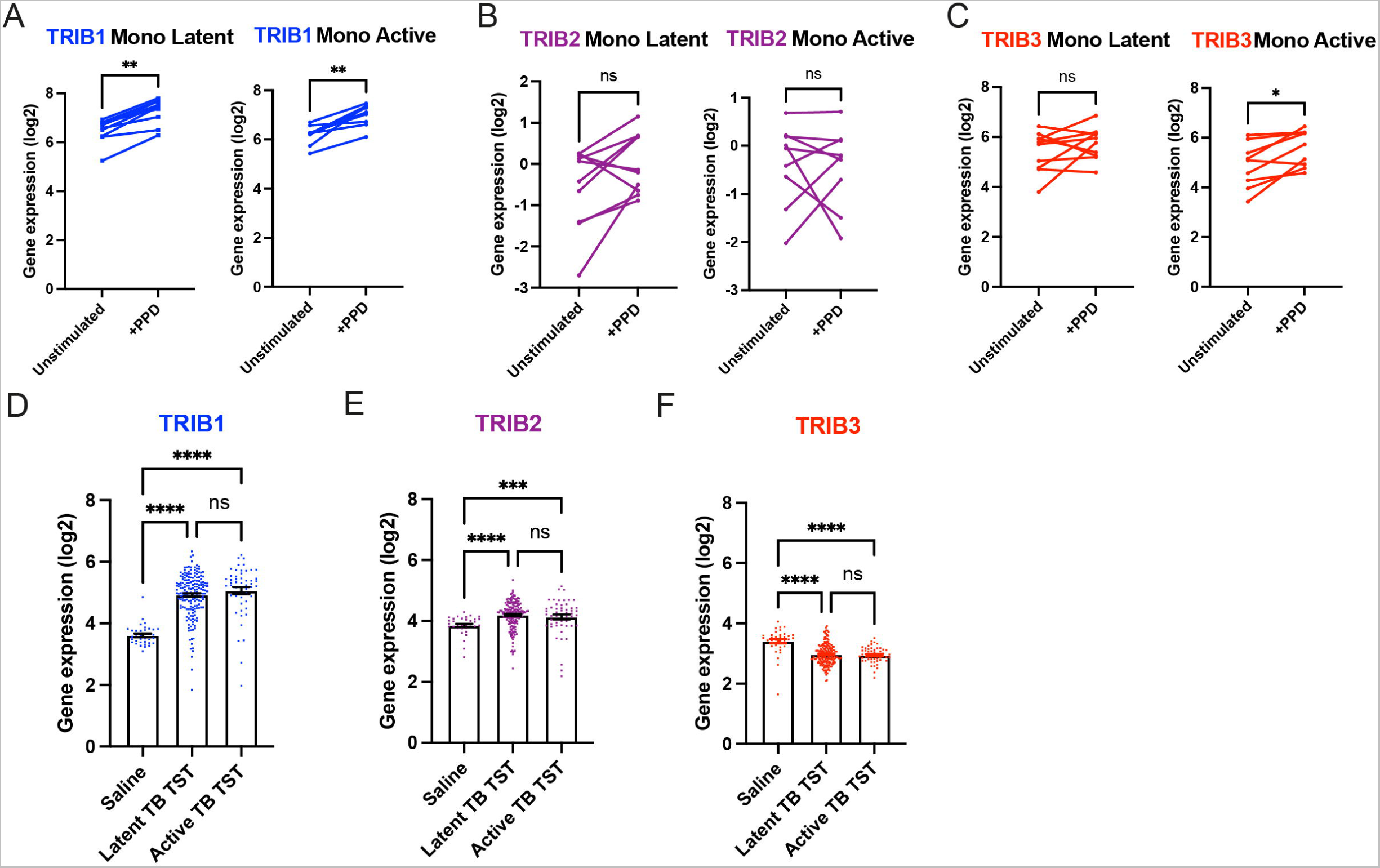
Expression of *TRIB1* in human monocytes and tissues is elevated after mycobacterial antigen stimulation. (A-C) Expression of *TRIB1, TRIB2 and TRIB3* transcripts in human CD14+ monocytes in patients with active or latent TB before and after 4 hours of *Mtb* protein derivative (PPD) stimulation *in vitro*. Each paired data point represents one individual, with active or latent TB (n = 9 and n = 7 respectively). Statistical significance determined by paired Wilcoxon tests P values shown are: \**P* < .05, \*\**P* < .01, \*\*\**P* < .001 and \*\*\*\**P* < .0001. (D-F) Expression of *TRIB1, TRIB2 and TRIB 3* within in saline injected human skin and from biopsies of the site of a tuberculin skin test (TST) in patients with active or latent TB. Each point represents one individual with bars for each group representing mean gene expression. n = 48 and 191 individuals with active or latent TB respectively. Statistical significance determined via Kruskal-Wallis with multiple comparisons. P values shown are: \**P* < .05, \*\**P* < .01, \*\*\**P* < .001 and \*\*\*\**P* < .0001.

Together these data reveal that TRIB1, and to a lesser extent TRIB2, expression is increased in response to both *in vitro* and *in vivo* mycobacterial antigen exposure in humans, independent of clinical TB disease grouping. We interpret these data as signifying a potential functional role for these pseudokinase in regulation of mycobacterial infections *in vivo*, but indicating the need for a tractable *in vivo* model of mycobacterial infection to study this further.

### Zebrafish Tribbles isoforms share homology with their human and mouse counterparts and are expressed in immune cell populations

To explore the functional role *in vivo* for Tribbles in the control of mycobacterial infections *in vivo*, we developed a zebrafish model of *Mycobacterium marinum* infection and *tribbles* manipulation.

Zebrafish have a single orthologue of each mammalian tribbles isoforms, with *tribbles1* (ENSDARG00000110963/previously ENSDARG00000076142), *tribbles2* (ENSDARG00000068179) and *tribbles3* (ENSDARG00000016200) genes. The exon organisation of Tribbles genes is conserved between human and zebrafish *trib* isoforms. Zebrafish *trib1* has 3 exons like murine *Trib1* and human *TRIB1* (Figure 2A). Human *TRIB2* and mouse *Trib2* also share this exon structure but are larger than the TRIB1 isoforms (Figure 2A). Zebrafish *trib2* is smaller than human *TRIB2* and mouse *Trib2* at 18.84kb and only possesses two coding exons (Figure 2A). Human *TRIB3*, mouse *Trib3* and zebrafish *trib3* share a similar exon organisation with a small non-coding first exon, followed by three coding exons (Figure 2A). Homology between Tribbles isoforms across species is not only observed at the genetic level, but also at the protein level (Hegedus et al. 2006). Tribbles have three key protein domains: an N terminal PEST domain, a pseudokinase domain and a functional C terminal (Hegedus et al. 2007). The pseudokinase contains a substrate binding site within its catalytic loop, and the functional C terminus contains two binding sites for either MEK or COP enzymes (Qi et al. 2006; Yokoyama et al. 2010). These three binding sites were compared across human, mice and zebrafish using the NCBI BLAST Global align online tool (Figure 2B). The pseudokinase catalytic loops in all three Tribbles family proteins (TRIB1-3), are found in human, mouse and zebrafish. In the case of TRIB1 and TRIB2 there is no variation in the amino acid sequence of the pseudokinase catalytic loop across the three species (Figure 2B). The pseudokinase catalytic loop of both mouse TRIB3 and zebrafish Trib3 differ slightly from Human TRIB3 with two amino acids that are different in mouse TRIB3 and one amino acid difference is observed in zebrafish Trib3. The amino acid sequences of human and zebrafish Tribbles were compared using the NCBI global align tool. Zebrafish Trib1 had the highest percentage identity when compared with human TRIB1 (52%), but also shared sequence homology with human TRIB2 with the highest identities match (66%) (Figure 2C). Zebrafish Trib2 shared the highest percentage identity with human TRIB2 (47% and 54% respectively). Zebrafish Trib3 had high identity matches for both human TRIB2 and TRIB3 (68% and 64% respectively) (Figure 2C). The overall size of Tribbles proteins remains consistent between human and mouse isoforms, with both human and mice TRIB1 sized at 372 amino acids (aa), human and mouse TRIB2 sized at 343aa. Mouse TRIB3 is 4aa shorter than human TRIB3 (354aa compared to 358aa). The zebrafish Tribbles isoforms are generally smaller proteins compared to the human and mouse Tribbles, with zebrafish Trib1 23aa smaller (at 349aa), Trib2 136aa smaller (at 207aa) and Trib3 10aa (at 348aa) compared to the human TRIB isoforms (Figure 2D).

**Figure 2:**
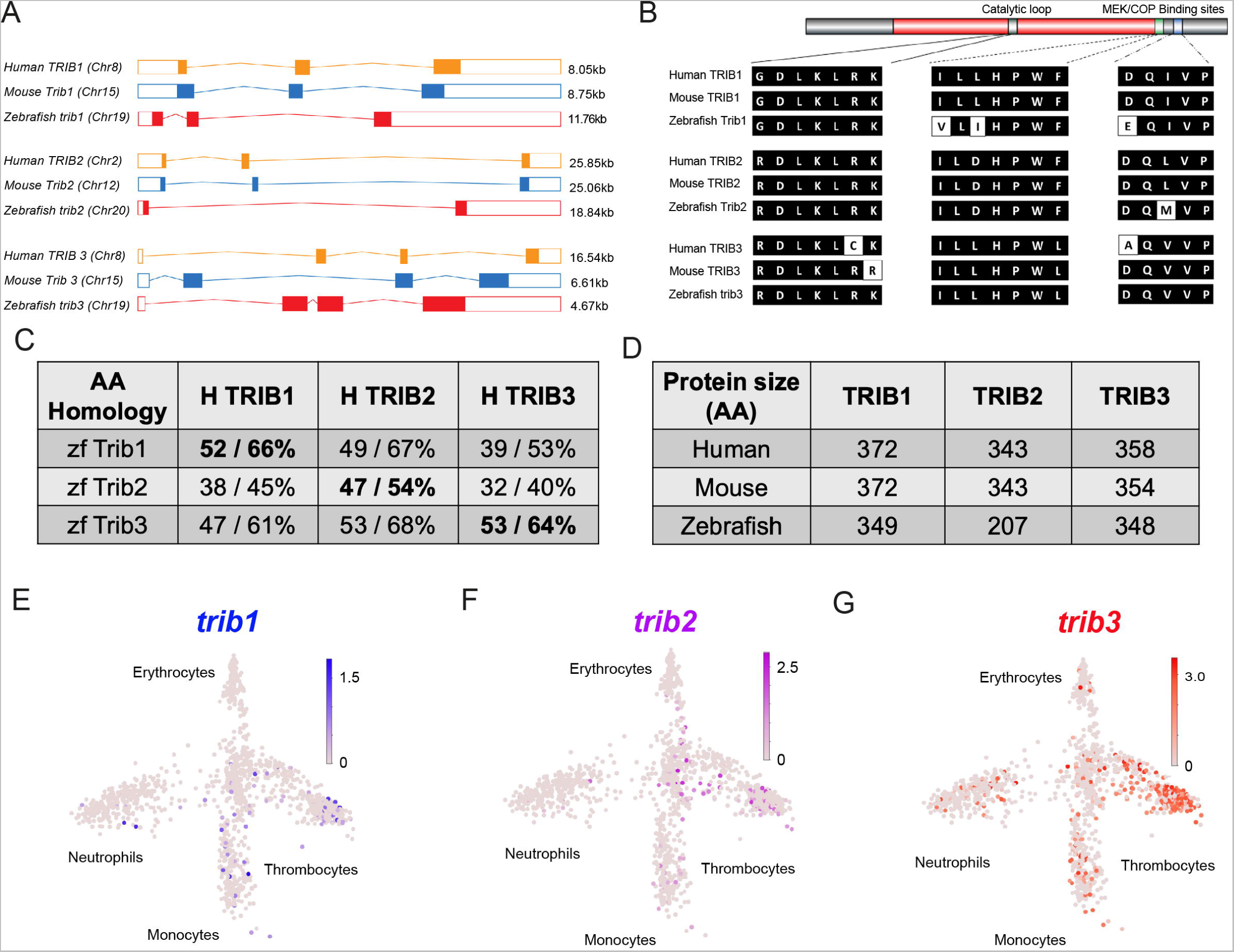
Zebrafish Tribbles share homology with their human and mice counterparts and are expressed in immune cell subpopulations. (A) The gene organisation of human (orange) TRIB1, mouse (blue) Trib1 and zebrafish (red). Exon maps produced from Ensembl database. Chromosome number location (chr) and transcript sizes in kilobases (kb) are shown. (B) Comparison of the three catalytic domains of Tribbles; the pseudokinase catalytic loop and MEK / COP1 bind sites, reveals high homology between species. (C) NCBI BLAST Global align revealed a high amino acid (AA) homology between zebrafish (zf) and human Tribbles protein sequences. Values described are positives / identities. (D) Protein sizes of the first and largest protein coding transcript of each gene are depicted in the number of AA, values obtained from Ensembl and Uniprot databases. (E-G) Gene expression of adult zebrafish leukocytes determined using the Zebrafish Blood Atlas (Athanasiadis et al. 2017). Each point represents a separate scRNAseq sample (cell); replicates performed across multiple zebrafish wildtype and transgenic strains. Each arm of schematic indicates A separate blood cell population (labelled). Deeper colour indicates higher expression (log10 Scale bars described for each gene).

To characterise the localisation of *trib* expression across the zebrafish larvae, whole mount *in situ* hybridisation probes were developed for each zebrafish *trib* isoform. All *tribbles* isoforms showed highest expression in the brain of the developing zebrafish larvae at 3dpf compared to sense probe controls (Figure S1). Expression of *trib* isoforms in immune cells was not detected by whole-mount *in situ* hybridisation of unchallenged larvae, compared to the expression of the highly expressed immune gene *l-plastin* (Figure S1). However, this does not negate low, sub-threshold, levels of *tribbles* isoforms in immune cells. *trib* levels in blood cell lineages were assessed using the Zebrafish Blood Atlas web tool ((Athanasiadis et al. 2017) https://scrnaseq.shinyapps.io/scRNAseq_blood_atlas/), based on scRNAseq of adult zebrafish leukocytes. All *trib* isoforms were expressed in subpopulations of neutrophils, monocytes and thrombocytes. *trib3* was expressed more abundantly and in a larger number of single cell RNAseq samples than other *trib* isoforms, and was found in macrophages, neutrophils and thrombocytes (Figure 1E-G).

In summary, zebrafish, mouse and human Tribbles share sequence similarity and have similar gene organisation and conserved catalytic binding sites, making zebrafish a viable model to explore a physiological role for human Tribbles in mycobacterial infections. Zebrafish express *trib* isoforms in immune cell subpopulations in resting conditions, suggestive of roles in regulating innate immunity.

### Overexpression of *trib1* is host-protective in a zebrafish mycobacteria infection model

To better understand how Tribbles can influence innate immunity and infection, genetic tools were generated to manipulate expression of zebrafish *trib* isoforms. Overexpression of zebrafish *tribbles* isoforms was achieved by injection of RNA at the one-cell stage. Injection of either *trib1*, *trib2* or *trib3* RNA did not grossly affect larval development, with embryos developing with no obvious adverse effects (Figure S2A-B). To determine outcomes in infection, a zebrafish *Mycobacterium marinum* (*Mm*) larval model was used, in which *<u>trib</u>* RNAs were injected at the one-cell stage, leading to ubiquitous overexpression. Overexpression of *trib1* significantly decreased bacterial burden of *Mm* by approximately 50% (p< 0.001) compared to the vehicle control, phenol red (PR) (Figure 3A-B). Dominant active *hif-1*α (DA1, an RNA shown to significantly reduce *Mm* burden by ∼50% (Elks et al. 2013) was used as a positive RNA control with dominant negative *hif-1*α (DN1, an RNA shown to have no significant effect on *M. marinum* burden) used as a negative RNA control (Elks et al. 2013). Overexpression of *trib2* also significantly reduced bacterial burden compared to the PR control, but not to the same extent as the positive DA *hif-1*α control nor *trib1* overexpression (Figure 3A-B). In contrast, overexpression of *trib3* had no significant effect on the levels of bacterial burden compared to the vehicle PR control (Figure 3A-B). Together, these data demonstrate that overexpression of *trib1* has the strongest host-protective effect compared to overexpression of other *trib* isoforms, reducing *M. marinum* burden by approximately 50%.

**Figure 3:**
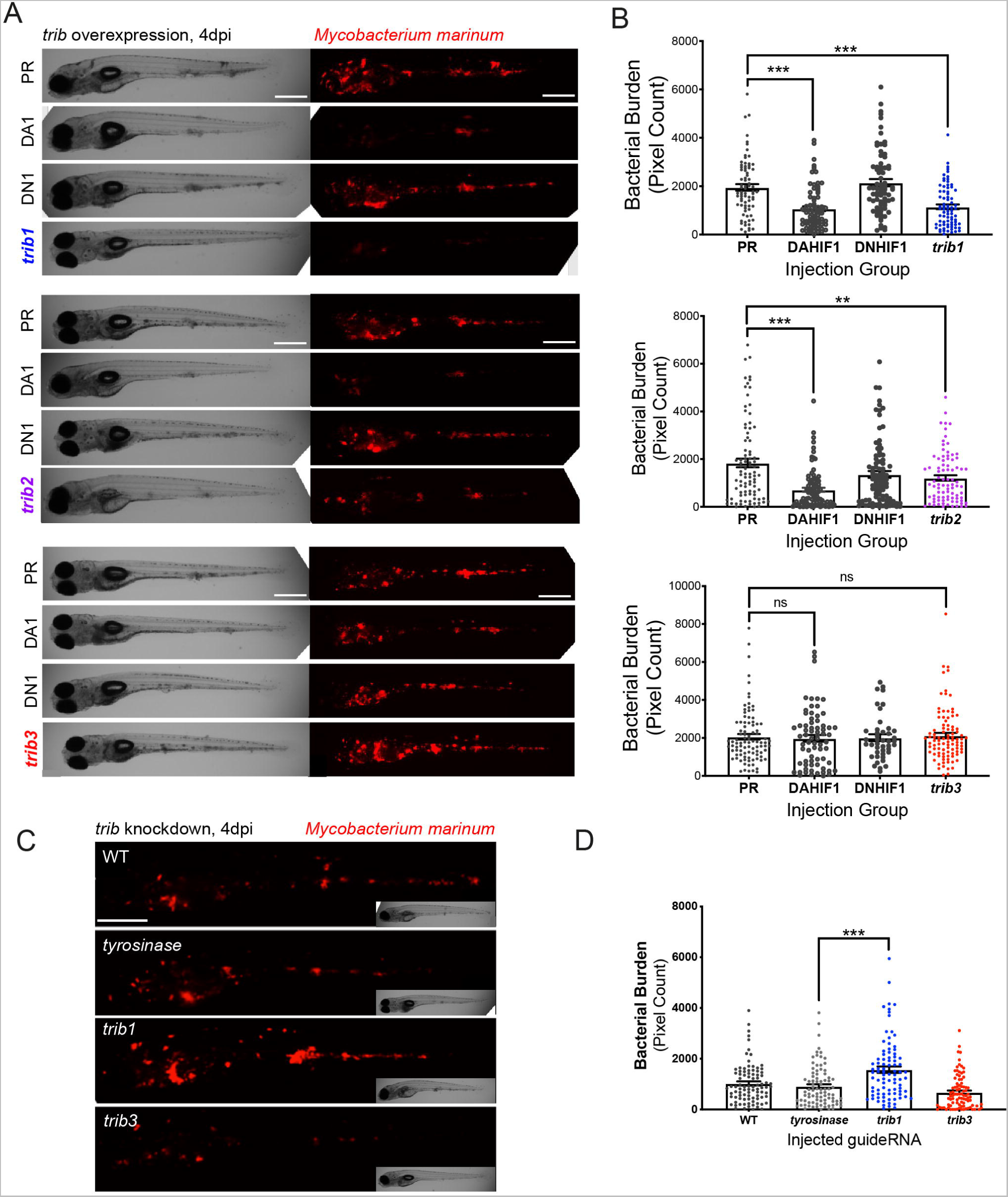
*trib1* overexpression is host protective against *Mm* infection. (A) Stereo-fluorescence micrographs of *Mm* mCherry infected 4dpi larvae after injection at the single-cell stage with DA Hif-1α (DA1), DN Hif-1 α (DN1), and *trib1, 2 and 3* using phenol red (control) as a negative vehicle control. DA1 and DN1 are RNA controls with DA1 having previously been shown to reduce infection levels. (B) Bacterial burden of larvae shown in (A). Data shown are mean ± SEM, n=76-77 in *trib1* experiment, n=86-89 in *trib 2* experiments and n=43-95 in *trib3* experiment, accumulated from 3 independent experiments for each *trib* gene. Statistical significance determined via one-way ANOVA with Bonferroni’s multiple comparisons. P values stated on graphs. (C) Stereo-fluorescence micrographs of Mm mCherry infected 4dpi larvae after injection with *tyrosinase* (control), *trib1* and *trib3* CRISPR guides (CRISPants). (D) Bacterial burden of larvae shown in (C). Data shown are mean ± SEM, n=87-90 fish accumulated from 3 independent experiments. Statistical significance determined via one-way ANOVA with Bonferroni’s multiple comparisons. P values shown are: \**P* < .05, \*\**P* < .01, \*\*\**P* < .001 and \*\*\*\**P* < .0001.

*trib* knockdown tools were developed using CRISPR-Cas9 technology. Guide-RNAs for each *trib* isoform were designed targeting the first coding exon of each *trib* gene and were injected into one-cell stage embryos, with *tyrosinase* (a control CRISP-ant which has negligible effects on innate immunity (Isles et al. 2019)) CRISP-ant as a negative control. CRISP-ant efficiency was tested using PCR and restriction enzyme digest, with successful CRISP-ants disrupting the restriction site. Efficient guide-RNAs were developed for both *trib1* and *trib3* (Figure S3), however guide-RNAs for *trib2* did not cause efficient knockdown. *Trib1* CRISP-ants had a higher burden of Mm compared to *tyrosinase* and *trib3* CRISP-ants (Figure 2C-D).

### *trib1* overexpression increases production of pro-inflammatory factors

TRIB1 has previously been shown to affect immune cell differentiation, with full-body *Trib1* deficient mice possessing a greater number of neutrophils and a reduced number of anti-inflammatory macrophages compared to wild-type (Satoh et al. 2013). Zebrafish *trib1* manipulation had the most profound effect on host pathogen interaction, with overexpression reducing Mm bacterial burden and CRISP-ant knockdown increasing burden. We therefore investigated the roles of *trib1* manipulation on the innate immune system.

Zebrafish *trib* isoforms were manipulated in neutrophil and macrophage transgenic reporter lines *Tg(mpx:GFP)i114* and *Tg(mpeg:nlsclover)sh436* and whole-body fluorescent cell counts were performed to assess whether *trib* manipulation influenced zebrafish leukocyte number. Neither *trib* overexpression nor CRISP-ant grossly affected neutrophil or macrophage numbers (Figure S4), suggesting that the host-protective effect of *trib1* overexpression is not due to an increase in number of innate immune cells. To investigate whether *trib1* influenced the inflammatory profiles of zebrafish leukocytes, production of the pro-inflammatory factors, *interleukin-1*β (*il-1*β) and nitric oxide (NO) were measured using a combination of transgenic reporter lines and immunostaining. Overexpression of *trib1* increased the levels of *il1*β*:GFP* (in a *Tg(il-1*β*:GFP)sh445* reporter line), to similar levels as the positive control DA Hif-1α, compared to phenol-red (PR) injected controls (Figure 4A-B). *trib3* overexpression did not increase levels of *il1*β*:GFP* and levels were similar to the negative controls DN Hif-1α and PR (Figure 4A-B). Similarly, *trib1* overexpression increased the levels of anti-nitrotyrosine staining, a proxy for immune cell antimicrobial nitric oxide production (Forlenza et al. 2008), to similar levels of DA Hif-1α (Elks et al. 2014; Elks et al. 2013) (Figure 4C-D). *trib3* overexpression did not increase levels of proinflammatory nitrotyrosine (Figure 4C-D).

**Figure 4:**
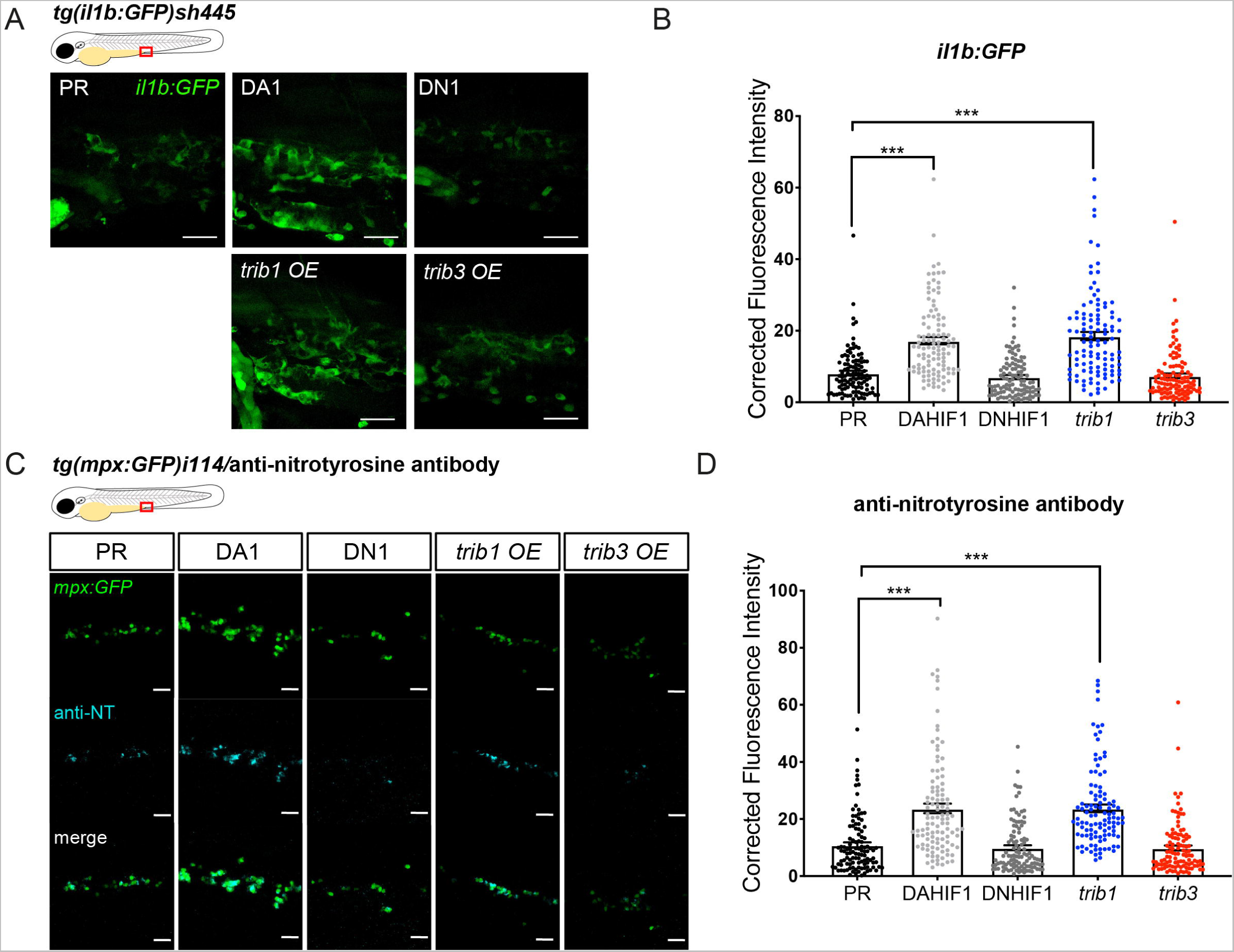
*trib1* overexpression increases production of proinflammatory *il-1*β and nitrotyrosine in the absence of infection. (A) Fluorescent confocal micrographs of 2dpf caudal vein region of *TgBAC(il-1β:eGFP)sh445* transgenic larvae. *il-1β:GFP* expression was detected by GFP levels. Larvae were injected at the 1 cell stage with dominant negative (DN) or dominant active (DA) Hif-1α or phenol red (PR) controls and *trib1* and *trib3* test RNAs. Scale bars = 25μm. (B) Corrected fluorescence intensity levels of *il-1β:GFP* confocal z-stacks in uninfected larvae at 2dpf of data shown in (A). Dominant active Hif-1α (DA1) controls and *trib1* fish had significantly increased *il-1β:GFP* levels in the absence of Mm bacterial challenge compared to phenol red (PR) and dominant negative Hif-1α (DN1) injected controls and *trib3* RNA injected embryos. Data shown are mean ± SEM, n=108 cells from 18 embryos accumulated from 3 independent experiments. Statistical significance was determined using one-way ANOVA with Bonferroni’s multiple comparisons post hoc test. P values shown are: \**P* < .05, \*\**P* < .01, \*\*\**P* < .001 and \*\*\*\**P* < .0001. (C) Fluorescence confocal z-stacks of the caudal vein region of 2dpf *mpx:GFP* larvae (neutrophils) immune-labelled with anti-nitrotyrosine (cyan) in the absence of Mm infection. Larvae were injected at the 1 cell stage with dominant negative (DN) or dominant active (DA) Hif-1α or phenol red (PR) controls and *trib1* and *trib3* test RNAs. Scale bars = 25μm. (D) Corrected fluorescence intensity levels of anti-nitrotyrosine antibody confocal z-stacks shown in (C). Data shown are mean ± SEM, n=108 cells from 18 embryos accumulated from 3 independent experiments. Statistical significance was determined using one-way ANOVA with Bonferroni’s multiple comparisons post hoc test. P values shown are: \**P* < .05, \*\**P* < .01, \*\*\**P* < .001 and \*\*\*\**P* < .0001.

### *Trib1*overexpression does not activate Hif signalling

Due to the protective effect of *trib1* overexpression closely mimicking that of DA-Hif1α a potential mechanistic link between the *hif-1*α and *trib1* pathways was investigated. *trib1* and *trib3* were overexpressed in a Hif-α transgenic reporter line, *Tg(phd3:GFP)i144* (*phd3* is a downstream target of Hif-α signalling) (Santhakumar et al. 2012). Neither *trib1* nor *trib3* overexpression activated the *phd3:GFP* line to detectable levels, indicating that *trib1* overexpression is not substantially increasing Hif-1α signalling to mediate *Mm* control (Figure 5). These data suggesting that the protective effects of *trib1* act via a different mechanism than Hif-1α activation.

**Figure 5:**
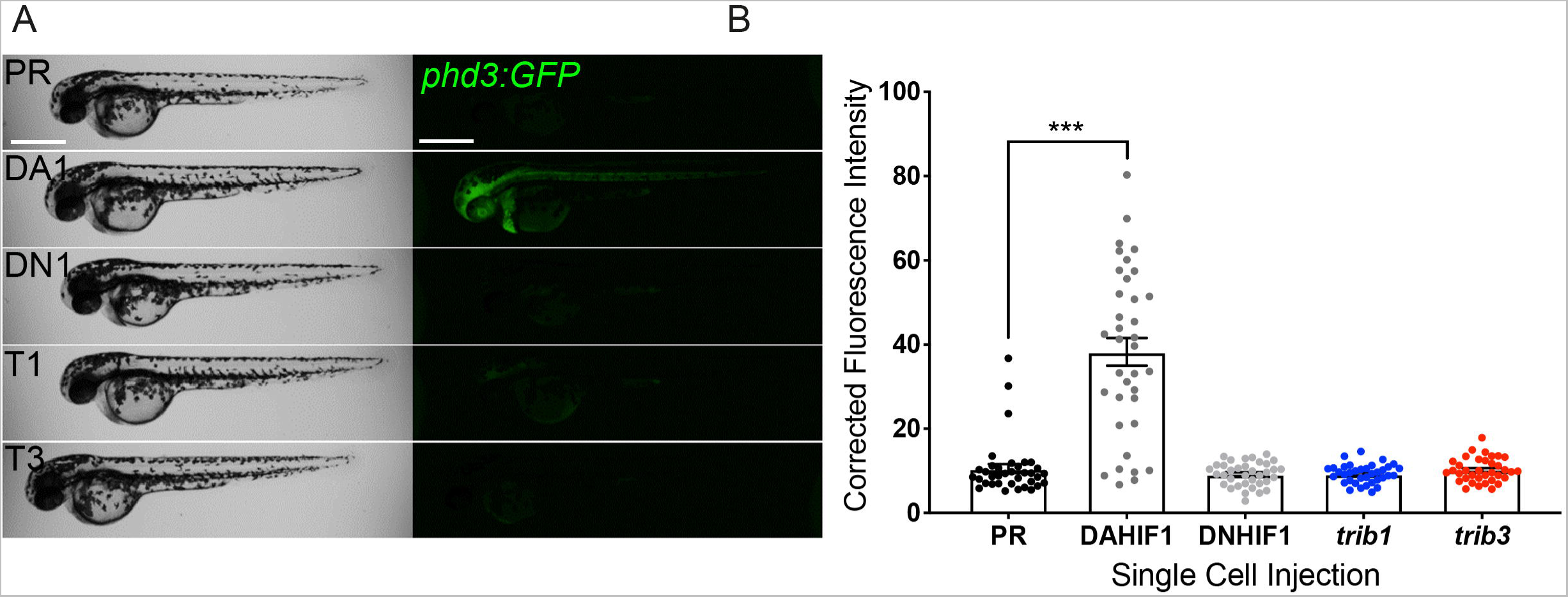
*trib1* and *trib3* overexpression do not induce expression of the Hif reporter *phd3:GFP*. (A) Stereo-fluorescence micrographs of 2dpf *phd3:GFP* larvae injected with phenol red (PR), DA Hif-1α and DN Hif-1α controls alongside *trib1* (T1) and *trib3* (T3) RNA. (B) Corrected fluorescence intensity levels of *phd3:GFP* larvae shown in (A). Only the DA Hif-1α injection led to increased Hif reporter levels compared to negative controls (PR and DN1) with *trib1* and *trib3* RNAs having no effect on *phd3:GFP* levels. Data shown is from 3 independent experiments, total 30 fish per group. Error bars depict SEM. Statistical significance determined through one-way ANOVA with multiple comparisons. P values shown are: \**P* < .05, \*\**P* < .01, \*\*\**P* < .001 and \*\*\*\**P* < .0001.

### The host protective effect of *trib1* is dependent on *cop1*

An important binding partner of the TRIB1 protein is the E3 ubiquitin ligase, COP1 (Jamieson et al. 2018; Kung and Jura 2019; Murphy et al. 2015). To investigate whether the host-protective effects of *trib1* overexpression in *Mm* infection were *cop1*-mediated, a *cop1* CRISPant was generated.

The zebrafish *cop1* gene (ENSDARG00000079329) is located on the forward strand of chromosome 2 and has 20 exons, all of which are coding (Figure S4). It has a single coding transcript, producing a Cop1 protein of 694 amino acids. The zebrafish *cop1* gene shares synteny and conserved sequence with both the human COP1 and murine Cop1 (determined using the ZFIN database (https://zfin.org/)).

In order to investigate whether the protective effect of *trib1* overexpression is *cop1*-mediated, *trib1* overexpression was combined with *cop1* CRISPants in *Mm* infected larvae. As previously observed, overexpression of *trib1* significantly reduced bacterial burden compared to phenol red controls when co-injected with *tyrosinase* guide (Figure 5A-B). The bacterial burden of *cop1* CRISPants, was not significantly different to the tyrosinase control group nor the tyrosinase control with *trib1* overexpression group. When *trib1* was overexpressed in *cop1* CRISPants, there was no significant decrease in burden, with the protective effect of *trib1* lost (Figure 6B) indicating that the protective effect of *trib1* overexpression is dependent on *cop1*.

**Figure 6:**
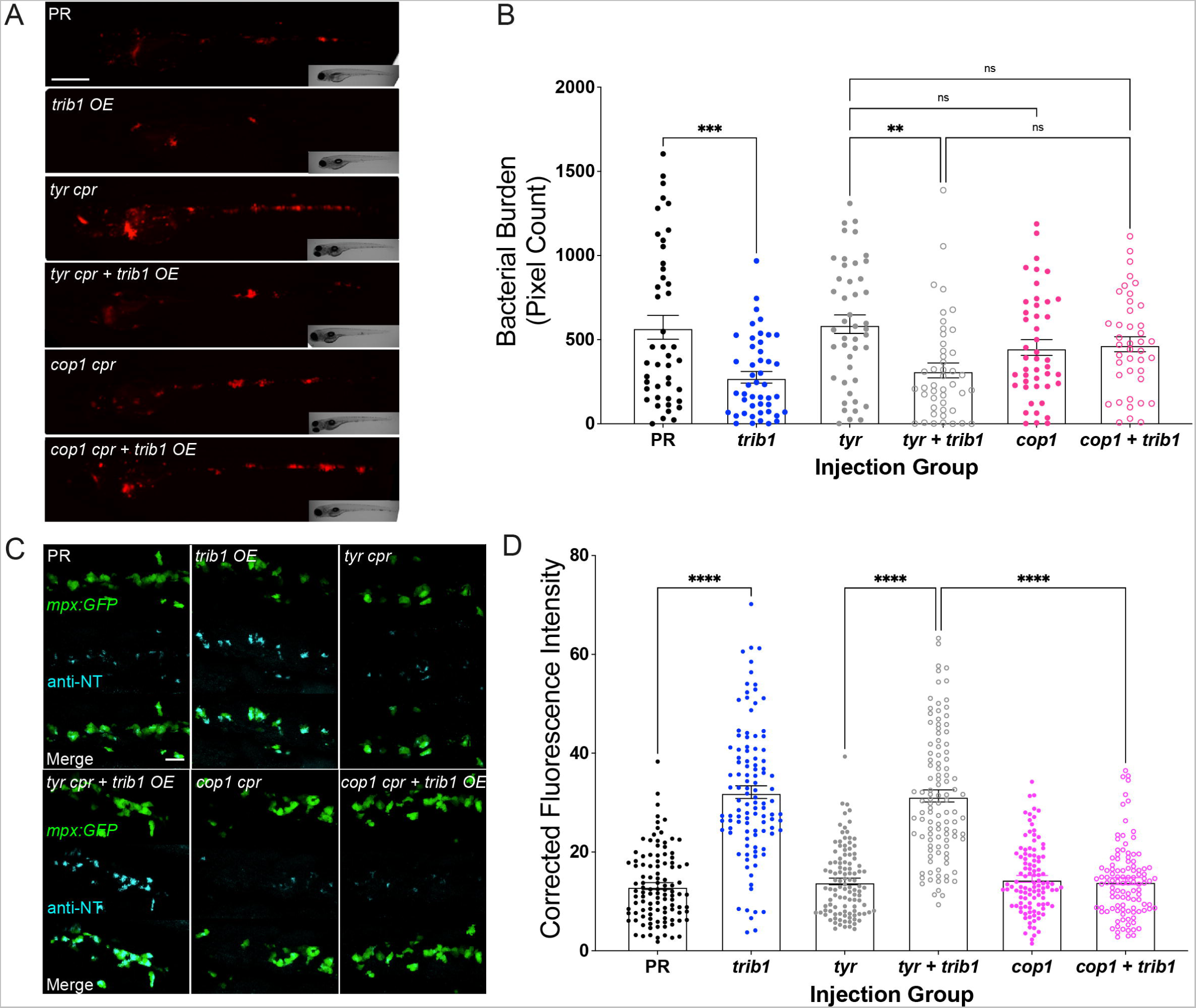
The host protective effect of *trib1* overexpression requires *cop1*. (A) Stereo-fluorescence micrographs of Mm mCherry infected 4dpi larvae after injection with *trib1* RNA (overexpression, OE) and *cop1 guide* RNA (CRISPants, cpr) using phenol red (vehicle) and tyrosinase (unrelated guide RNA) CRISPants as negative controls. (B) Bacterial burden of larvae shown in (A). Data shown are mean ± SEM, n=71-76 accumulated from 3 independent experiments. Statistical significance determined via one-way ANOVA with Bonferroni’s multiple comparisons. P values shown are: \**P* < .05, \*\**P* < .01, \*\*\**P* < .001 and \*\*\*\**P* < .0001. (C) Fluorescence confocal z-stacks of the caudal vein region of 2dpf *mpx:GFP* larvae (neutrophils) immune-labelled with anti-nitrotyrosine (cyan) in the absence of Mm infection. Larvae were injected at the 1 cell stage with *trib1* RNA (overexpression, OE) and *cop1 guide* RNA (CRISPants, cpr) using phenol red (vehicle) and tyrosinase (unrelated guide RNA) CRISPants as negative controls. Scale bars = 25μm. (D) Corrected fluorescence intensity levels of anti-nitrotyrosine antibody confocal z-stacks shown in (C). Data shown are mean ± SEM, n=108 cells from 18 embryos accumulated from 3 independent experiments. Statistical significance was determined using one-way ANOVA with Bonferroni’s multiple comparisons post hoc test. \**P* < .05, \*\**P* < .01, \*\*\**P* < .001 and \*\*\*\**P* < .0001.

The effect of *cop1* knockdown on the production of antimicrobial NO production was investigated using the anti-nitrotyrosine antibody. Overexpression of *trib1* significantly increased neutrophil anti-nitrotyrosine fluorescence levels compared to the PR control in the *tyrosinase* controls (Figure 6C-D). The *cop1* CRISPant group possessed comparable anti-nitrotyrosine levels to both the PR and tyrosinase control groups. *trib1* overexpression in the *cop1* CRISPants did not increase anti-nitrotyrosine levels and instead was comparable with the *cop1* CRISPants alone and both PR and tyrosinase controls (Figure 6D).

Together these data show that when *cop1* is knocked down the antimicrobial and host protective effects of *trib1* overexpression are lost, indicating a dependency of the *trib1* effect on *cop1*.

## Discussion

TRIB1 has previously been shown to be a key regulator of multiple inflammatory factors and inflammatory cell function, influencing pathologies with an inflammatory component including cancer and atherosclerosis (Johnston et al. 2015). Innate immunity and production of inflammatory factors are key defence mechanisms against invading pathogens, yet the role of Tribbles in the immune response to infection is poorly understood. We provide evidence that mycobacterial antigen stimulation both *in vitro* and *in vivo* induces human TRIB1 and TRIB2 expression independent of TB disease status. We used the zebrafish TB model to show that this expression has key and newly appreciated functional roles. *trib1* is required to control Mm infection *in vivo*, associated with increase production of antimicrobial factors, such as *il-1*β and NO. We also show a role for *cop1*, a key binding partner of TRIB1, which is required for the host-protective effects of *trib1* overexpression. The novel *in vivo* tools developed to investigate the immune roles of tribbles in zebrafish, create new opportunities to further investigate Tribbles 1 as a potential therapeutic target, not only in infection, but in a wider range of disease contexts that have an innate immunity component.

For the first time we have identified that Tribbles 1 is an important isoform in the host response to mycobacterial infection, with *TRIB1* being upregulated in human monocytes after mycobacterial antigen challenge and early overexpression of *trib1* being host-protective in a zebrafish TB model. This is in line with literature showing that TRIB1 has a key role in the regulation of pro-inflammatory profiles (Arndt et al. 2018; Niespolo et al. 2020; Ostertag et al. 2010). If inflammatory signals are initiated early in infection, this can improve infection outcomes and reduce bacterial burden, whereas later induction of these signals could be harmful to the host. An example of this is control of inflammatory response with HIF-1α signalling, where early stabilisation and activation of HIF-1α signalling is beneficial (Elks et al. 2013; Lewis and Elks 2019; Ogryzko et al. 2019), but late activation, or excessive HIF-1α is hyper-inflammatory and can increase bacterial burden in animal models (Braverman and Stanley 2017; Domingo-Gonzalez et al. 2017).

TRIB1 is a well-known regulator of innate immune cells and functions. *Trib1-/-* mice have a defective inflammatory response, with reduced pro-inflammatory gene expression (including Nos2 and Il-1β compared to controls) resulting in a defective pro-inflammatory macrophage response, with BMDMs producing less NO and defective phagocytosis (Arndt et al. 2018). In zebrafish, *trib1* overexpression increased production of pro-inflammatory factors, indicating this control of inflammatory factors NO and Il-1β via TRIB1 is conserved in fish. The NO response generated by TRIB1 may be produced through JAK/STAT signalling, which TRIB1 regulates to influence macrophage polarisation phenotypes via STAT3 and STAT6 (Arndt et al. 2018). Polarised macrophage subsets have also been identified in the zebrafish model, with heterogeneity observed in the macrophage population with inflammatory markers (Hammond et al. 2023; Nguyen-Chi et al. 2015). It is unclear whether zebrafish *trib1* could regulate macrophage inflammatory profiles via STAT3 and STAT6 as in the murine model. However, as zebrafish Stat3 has roles in macrophage efferocytosis, survival and cytokine secretion (Campana et al. 2018) and Stat6 has roles in type 2 immune signalling (Cronan et al. 2021) this could be conserved and a potential mechanism of *trib1* regulation of inflammatory phenotypes.

TRIB1 has been associated with inflammation and immune response, whereas TRIB3 is strongly associated with metabolic function, including the regulation of glucose homeostasis (Angyal and Kiss-Toth 2012; Prudente et al. 2012; Zhang et al. 2013) which also has regulatory roles in innate immune cells such as macrophages (Steverson et al. 2016; Wang et al. 2012). There are robust links between glucose metabolism and innate immune responses, such as the glycolytic switch which is closely related to macrophage polarisation (Zhu et al. 2015). Both glucose and lipid metabolism have roles in infection defence when utilised by immune cells. In *Mtb* infection, lipid droplets produced by macrophages can be used as an antimicrobial mechanism (Knight et al. 2018), or a source of lipids for *Mtb* to utilise (Daniel et al. 2011) as a method to potentially manipulate host macrophage defence (Menon et al. 2019). This process could potentially be influenced by Tribbles. In murine atheroma models, Trib1 increased the lipid accumulation in macrophages leading to the formation of foam-cells (Johnston et al. 2019). In *Drosophila melanogaster*, *trbl* knockdown increased circulating triglyceride levels (Das et al. 2014) and in mice where TRIB3 knockdown in a murine adipose cell line (3T3-L1) increased intracellular triglycerides (Takahashi et al. 2008) and targeted deletion of murine TRIB3 resulted in elevated triglyceride levels in the liver (Örd et al. 2018).

It is interesting to note that overexpression of *trib* genes did not affect macrophage and neutrophil numbers in the zebrafish larvae, unlike in Trib1 deficient mice, where the number of neutrophils is increased due to dysregulated C/EBPα (Satoh et al. 2013). Similar to mammalian neutrophil differentiation, zebrafish neutrophil differentiation is partly regulated via C/EBP transcription factors including Cebpα (Dai et al. 2016), Cebp1 (the functional homolog of mammalian C/EBPε, (Kim et al. 2016)) and Cebpβ (Wei et al. 2020). It is therefore unclear why *trib1* manipulation did not affect neutrophil differentiation in the zebrafish and highlights potential differences between the function of zebrafish *trib1* compared to murine TRIB1.

The host protective effect of *trib1* overexpression closely mimicked the effects of Hif-1α stabilisation, with an increase in the production of anti-microbial factors NO and Il-1β and a decrease in *Mm* infection burden (Elks et al. 2013). HIF transcription factors respond to oxygen tension and are stabilised under hypoxic conditions. TRIB3 has been associated with HIF-1α in renal cell carcinoma patients and HIF-1α binds to multiple regions in the TRIB3 promoter, with HIF-1α overexpression resulting in upregulation of TRIB3 expression (Hong et al. 2019). In lung adenocarcinoma cells, TRIB3 knockdown decreased levels of HIF-1α (Xing et al. 2020), indicating a feedback loop between TRIB3 and HIF-1α, where one can regulate the other and vice-versa. In common with TRIB3, a potential link between TRIB2 and HIF-1α has been reported, as depletion of TRIB2 significantly decreased the effect of TNFα on HIF-1α stability and accumulation in multiple cancer cell lines (Schoolmeesters et al. 2012). However, there is no current link identified between TRIB1 and HIF-1α and this was reflected in our data showing that *trib1* overexpression did not lead to an increase in a well-validated Hif-α reporter line (Santhakumar et al. 2012).

Many of the reported regulatory functions of TRIB1 are dependent on COP1. As the protective effect of *trib1* overexpression was reduced when *cop1* was depleted, it appears there is some dependency on *cop1* expression to produce the protective effect of improving the host response to infection. Interestingly, *cop1* CRISPants possess slightly reduced burden compared to controls without the overexpression of *trib1*, suggesting that *cop1* depletion alone may offer a small level of protection. In cancer cell lines infected with *Mycobacterium bovis* Bacillus Calmette-Guérin (BCG), BCG induced Sonic Hedgehog signalling increasing COP1 expression, leading to the inhibition of apoptosis in the cell line (Holla et al. 2014), indicating there may be a COP1 response to mycobacterial infection.

Together our findings show a potential therapeutic application of targeting Trib1 to improve infection outcomes. It appears to control multiple pathways, we have demonstrated here *il-1b* and NO control, therefore it may be more effective than targeting one of these alone. Due to its potential functions in multiple pathways, any targeting of Trib1 must be carefully controlled. For example, overexpression of TRIB1 in chronic mycobacterial infections may be beneficial against infection, but could trigger immunopathology. The concept of host immunomodulation is an emerging therapeutic avenue for infectious disease, especially with the continually increasing problem of anti-microbial resistance in multiple pathogens, and could potentially be used alongside anti-microbial drug treatment. To aid the efficiency of host immunomodulation, and to help avoid off-target effects, specific targeting methods can be used. Polymersomes have been shown to be a promising avenue for drug delivery to immune cells and could be utilised for the delivery of host immunomodulatory compounds and factors (Fenaroli et al. 2020). Therefore, with targeted delivery methods and transient manipulation of TRIB1 through pharmacological or genetic approaches, this could potentially improve infection outcome of mycobacterial infection and pave the way for further research into TRIB1 as a target for host-derived therapies.

## Acknowledgements

The authors would like to thank Dr Heba Ismail, The University of Sheffield, for her expertise and helpful advice on E3 ubiquitin ligases. Thanks also to The Biological Services Aquarium Team at the University of Sheffield for their expert assistance with zebrafish husbandry.

