## Supplemental Figures and Legends for "Tribbles1 and Cop1 cooperate to protect the host during *in vivo* mycobacterial infection"

### Supplementary Figures and Legends

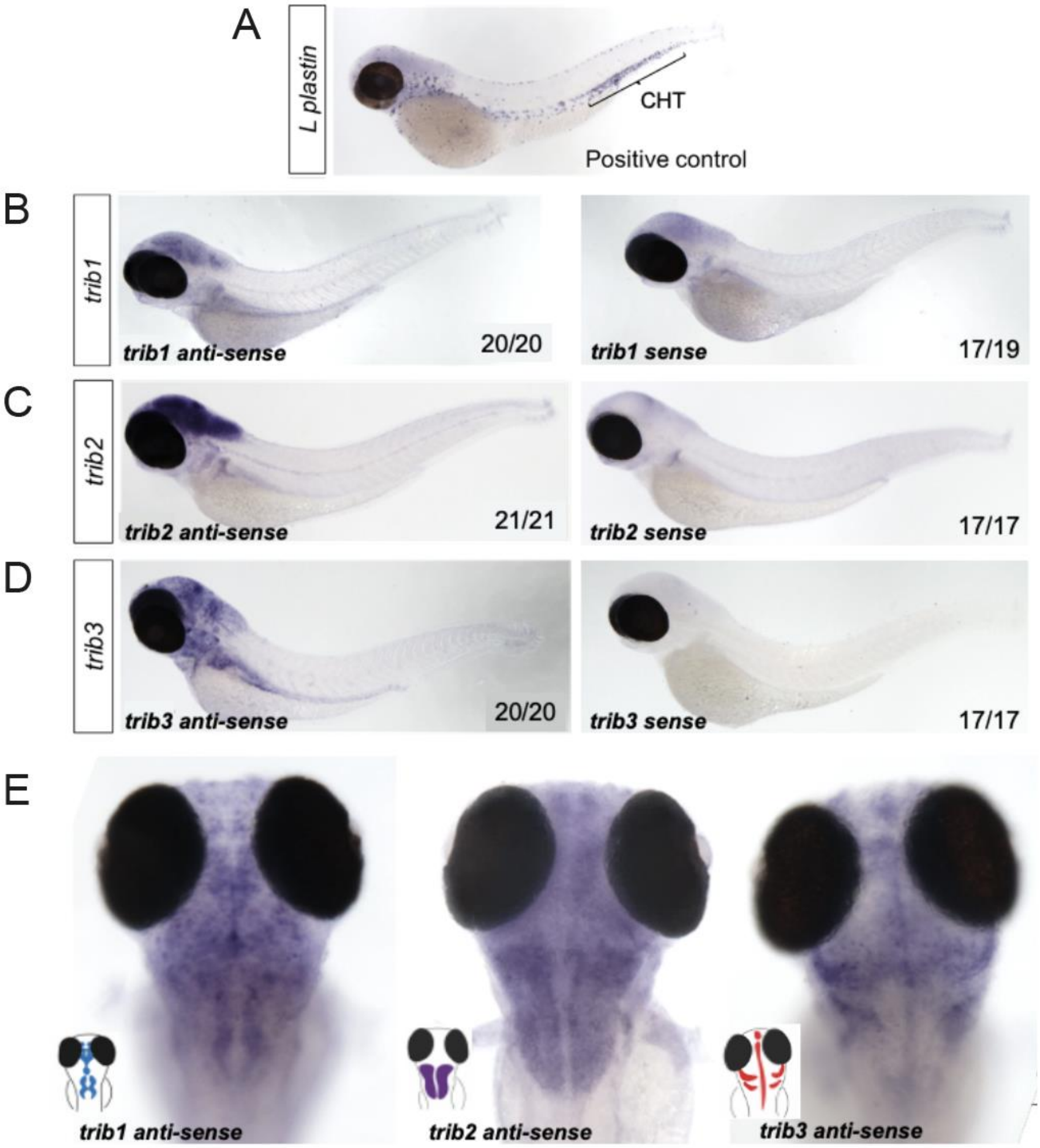


**Figure S1: Expression of zebrafish tribbles is located primarily in the brain, with each *trib* isoform displaying a distinct pattern of expression**

(A) Wholemount in situ hybridisation of the pan-leukocyte marker *L-plastin* was used as a positive control to stain leukocytes, present predominantly in the caudal hematopoietic tissue (CHT).

(B) Antisense and sense (control) wholemount *in situ* hybridisation of *tribbles1*. Numbers correspond to number of fish with pictured phenotype / total fish imaged. N=17-21 fish imaged per group from one independent experiment per *trib* isoform. Images are representative of phenotype from experimental groups. Imaged using 4x magnification using stereoscope and colour camera.

(C) Antisense and sense (control) wholemount in situ hybridisation of *tribbles1*.

(D) Antisense and sense (control) wholemount in situ hybridisation of *tribbles1*.

(E) Magnified dorsal views (8x magnification) showing distinct expression of each *trib* isoform in the brain (representative images from each group shown). Scale bars = 200µm.

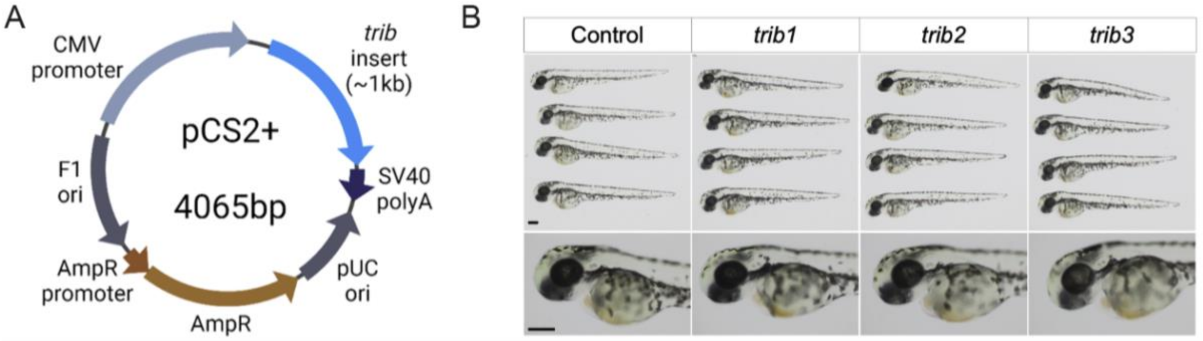


**Figure S2: tribbles RNA overexpression caused no observable developmental defects**

(A) cDNA sequences of zebrafish *trib* isoforms (*trib1*, *trib2* and *trib3*) were cloned into the expression plasmid pCS2+ .

(B) mRNA of *trib* isoforms (100ng/µl) was synthesised from pCS2+ plasmids and 1nl was injected into one-cell stage zebrafish embryos, which resulted in healthy development. Larvae imaged at 2dpf using stereoscope at 4x magnification (head images at 8x magnification). Scale bars = 200µM.

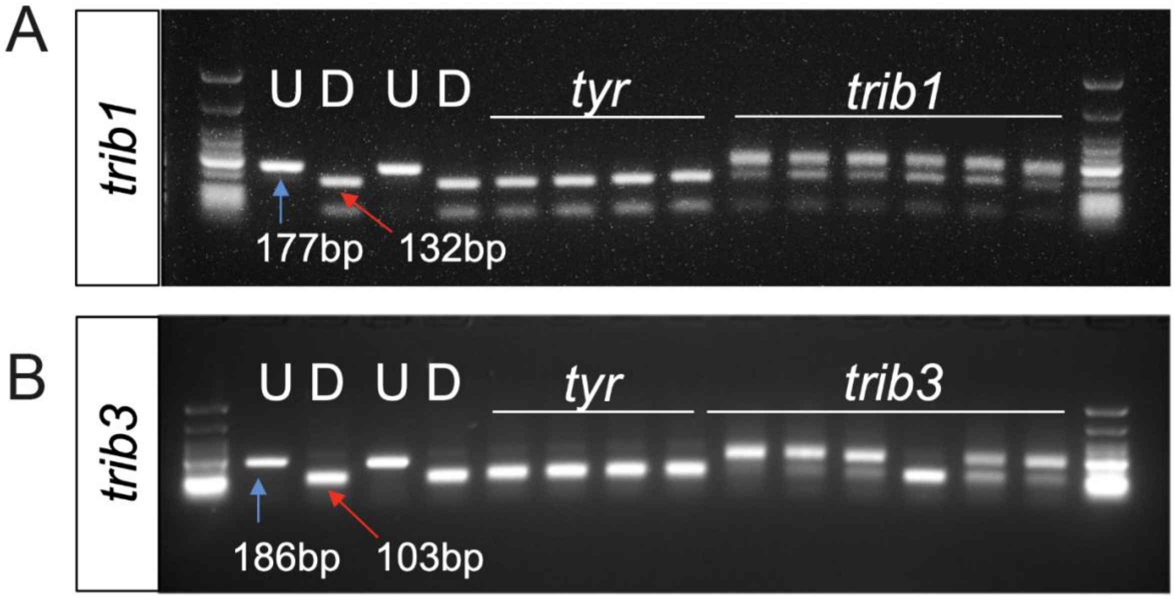


**Figure S3: GuideRNAs against trib1 and trib3 efficiently disrupt a restriction site.**

(A) Electrophoresis gel of larvae injected with *trib1* guide RNA with a SacII restriction digest. The undigested PCR band is 177bp and the successfully digested band is 132bp. Two uninjected fish were used as controls without (U) and with (D) digestion. Four *tyrosinase* (*tyr*) guide injected larvae had conserved SacII sites while 6/6 *trib1* larvae did not digest with SacII indicating a genomic disruption at the CRISPR PAM site. First and last lanes of gel images contain NEB Low Molecular Weight ladder for reference.

(B) Electrophoresis gel of larvae injected with *trib3* guide RNA with a MwoI restriction digest. The undigested PCR band is 186bp and the successfully digested band is 103bp. Two uninjected fish were used as controls without (U) and with (D) digestion. Four *tyrosinase* (*tyr*) guide injected larvae had conserved MwoI sites while 5/6 *trib3* larvae did not digest with MwoI indicating a genomic disruption at the CRISPR PAM site. First and last lanes of gel images contain NEB Low Molecular Weight ladder for reference.

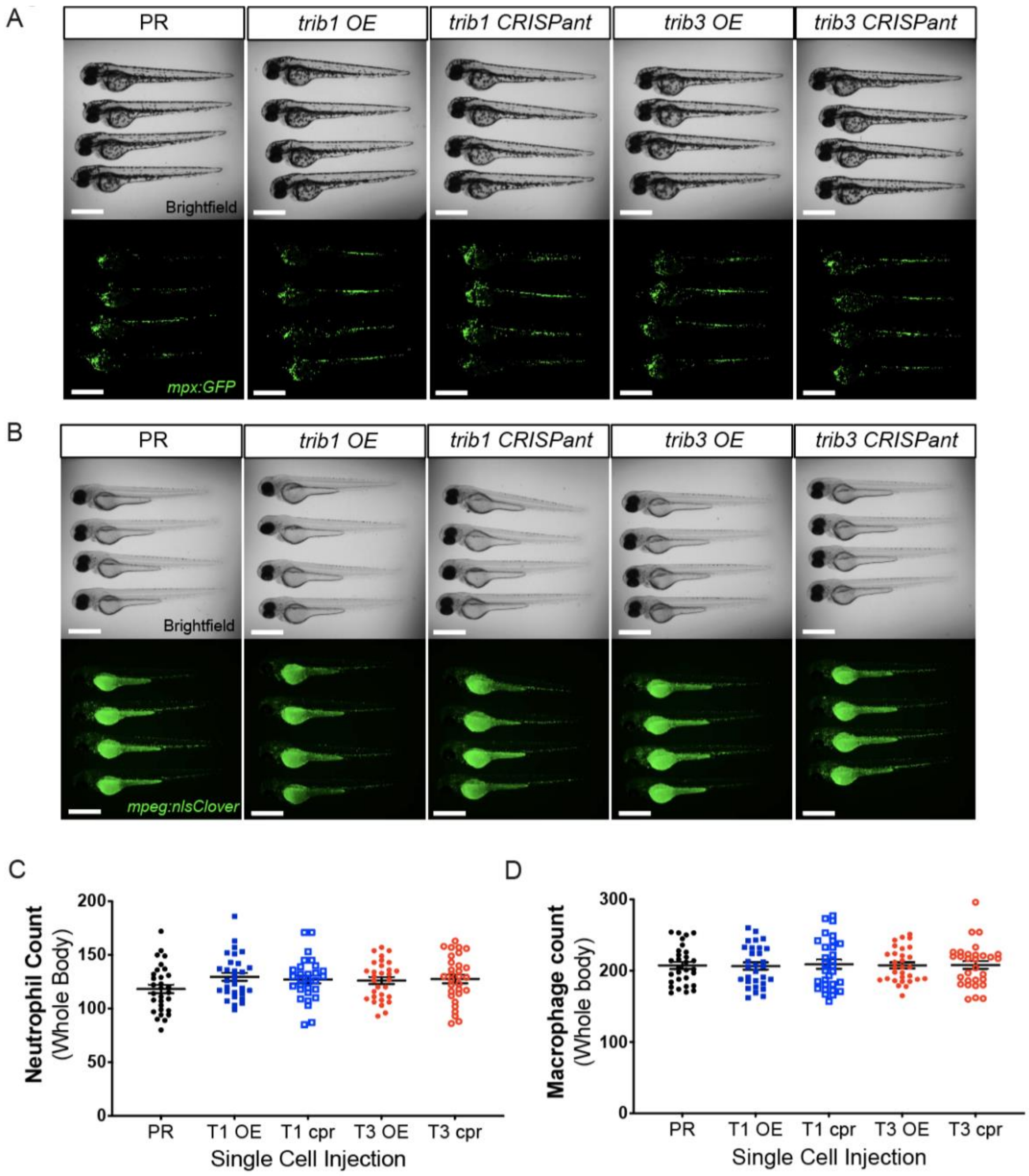


**Figure S4: Modulation of either *trib1* or *trib3* does not alter the number of macrophages and neutrophils in zebrafish larvae.**

Transgenic reporter lines for either neutrophils (Tg(mpx:GFP)i114, (A)) or macrophages (Tg(mpeg:nlsclover)sh436, (B)) were used for whole body cell counts of leukocytes. For overexpression (OE, full coloured symbols), 1nl of trib1 mRNA (100ng/µl), trib3 mRNA (100ng/µl), or phenol red (PR, diluted 1:10 in nuclease free water) vehicle control was injected into the yolk of single cell stage transgenic embryos. For knockdown, CRISPants (cpr, empty coloured symbols) were created through injecting 1nl of CRISPR mixes consisting of 1µl 20µM guideRNAs (for trib1 or trib3), 1µl Cas9 nuclease (diluted 1:3 in diluent B), 1µl 20µM tracrRNA (20µM) and 1µl DEPC water, was injected into the yolk of single cell stage transgenic embryos. At 2 days post fertilisation groups were first blinded, before fluorescent cells were counted by eye at a fluorescent stereomicroscope. Data are from 3 independent experiments, total 30 fish per group. Representative images from each group shown in A and B, scale bars = 1mm. Error bars (C-D) depict SEM. No significant differences observed between groups determined by one-way ANOVA with Bonferroni's multiple comparisons post hoc test.

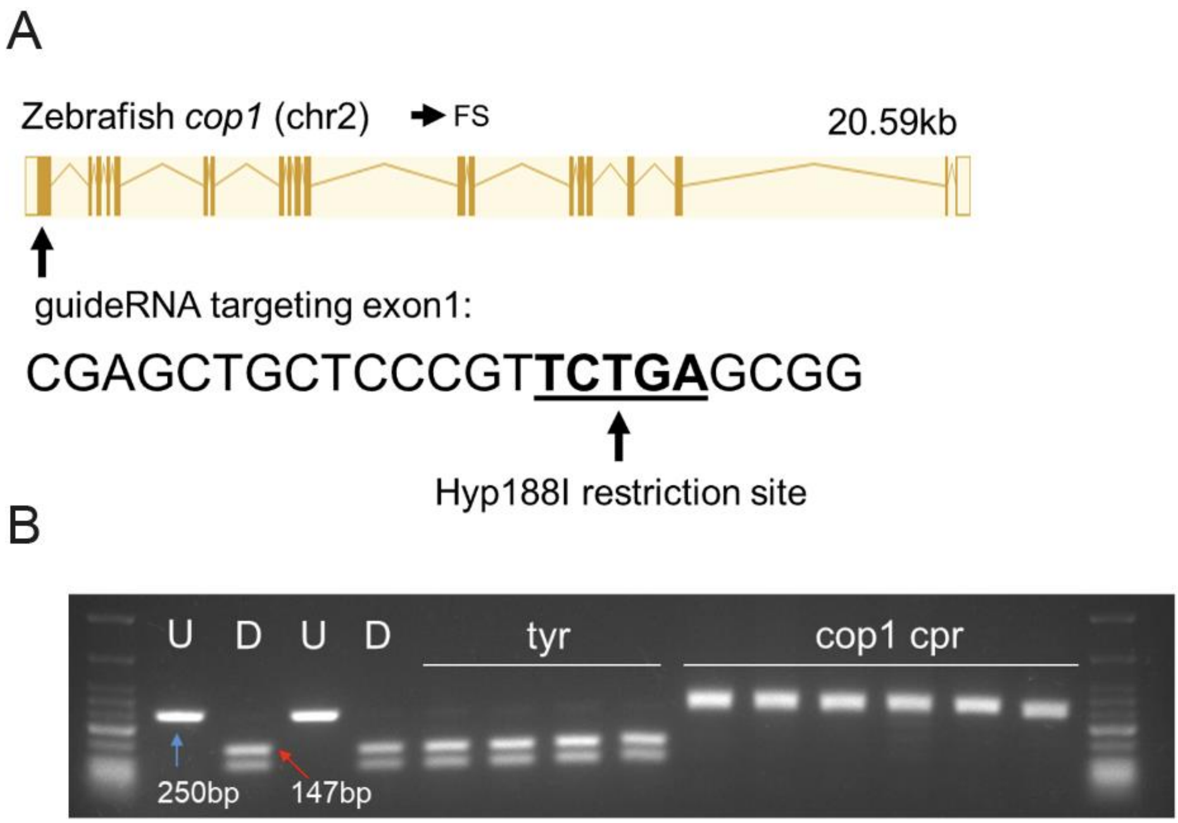


**Figure S5: An efficient CRISPR-Cas9 guideRNA was generated for the zebrafish cop1 gene.**

(A) The zebrafish cop1 gene (ENSDARG00000079329) has one coding transcript which was used as a basis for CRISPR-Cas9 guide design. Exon map and gene information obtained from Ensembl database. The web tool ChopChop was used to design a cop1 guide RNA with suitable restriction enzyme bind site (black line). PAM cut site (NGG) denoted with blue line.

(B) Electrophoresis gel of larvae injected with *cop1* guide RNA (cpr) with a Hyp188I restriction digest. The undigested PCR band is 250bp and the successfully digested band is 147bp. Two uninjected fish were used as controls without (U) and with (D) digestion. Four *tyrosinase* (*tyr*) guide injected larvae had conserved SacII sites while 6/6 *cop1* larvae did not digest with Hyp188I indicating a genomic disruption at the CRISPR PAM site. First and last lanes of gel images contain NEB Low Molecular Weight ladder for reference.
